# Toward subtask decomposition-based learning and benchmarking for genetic perturbation outcome prediction and beyond

**DOI:** 10.1101/2024.01.17.576034

**Authors:** Yicheng Gao, Zhiting Wei, Kejing Dong, Jingya Yang, Guohui Chuai, Qi Liu

## Abstract

Deciphering cellular responses to genetic perturbations is fundamental for a wide array of biomedical applications, ranging from uncovering gene roles and interactions to unraveling effective therapeutics. Accurately predicting the transcriptional outcomes of genetic perturbations is indispensable for optimizing experimental perturbations and deciphering cellular response mechanisms; however, three scenarios present principal challenges, i.e., predicting single genetic perturbation outcomes, predicting multiple genetic perturbation outcomes and predicting genetic outcomes across cell lines. In this study, we introduce <u>S</u>ub<u>TA</u>sk decomposition <u>M</u>odeling for genetic <u>P</u>erturbation prediction (STAMP), a conceptually novel computational strategy for genetic perturbation outcome prediction and downstream applications. STAMP innovatively formulates genetic perturbation prediction as a subtask decomposition (STD) problem by resolving three progressive subtasks in a divide-and-conquer manner, i.e., identifying differentially expressed gene (DEG) postperturbations, determining the regulatory directions of DEGs and finally estimating the magnitudes of gene expression changes. In addition to facilitating perturbation prediction, STAMP also serves as a robust and generalizable benchmark guide for evaluating various genetic perturbation prediction models. As a result, STAMP exhibits a substantial improvement in terms of its genetic perturbation prediction ability over the existing approaches on three subtasks and beyond, including revealing the ability to identify key regulatory genes and pathways on small samples and to reveal precise genetic interactions. Overall, STAMP serves as a fundamentally novel and effective prediction and generalizable benchmarking strategy that can facilitate genetic perturbation prediction, guide the design of perturbation experiments, and broaden the understanding of perturbation mechanisms.

## Main

The transcriptional outcomes of a single-cell genetic perturbation reveal fundamental insights into cell functions and how a cell responds to external interventions. Various high-throughput sequencing technologies, including single-cell RNA sequencing^1^ (scRNA-seq) and single-cell Clustered Regularly Interspaced Short Palindromic Repeats (CRISPR) genetic screening methods^2^ (for example, Perturb-seq^3^), have enabled the dissection of the transcriptional responses of individual cells to specific genetic perturbations, facilitating the uncovering of the roles and interactions of genes^4^. This has profound implications for investigating the regulatory mechanisms of genes and unraveling effective therapeutics in biomedical studies^5-7^. With recent advancements in experimental methodologies, high-throughput sequencing can now generate genetic perturbation outcomes for either a single gene or multiple genes in a cell^3, 8, 9^. However, because of the vast combinatorial explosion exhibited by the potential gene combination space, a brute-force experimental exploration of the outcome of perturbing multiple genes is impractical^10-12^. Furthermore, as single-cell CRISPR-based genetic screening technology is still in its early stage and cost-ineffective, limited cell line perturbation data have been obtained. Therefore, there is an urgent need to develop computational approaches that can guide the prediction of perturbed transcriptional outcomes in various scenarios involving single-gene perturbations, multiple-gene perturbations and cross-cell line adaptations.

Several tools have been developed for modeling in silico gene perturbations and predicting transcriptional genetic perturbation outcomes^6, 11, 13-18^. These tools are mainly categorized into three groups. (1) Inferring gene regulatory networks^6, 13^: Methods in this category focus on modeling in silico gene perturbations by inferring gene regulatory networks from both specialized and publicly available datasets, including CellOracle^6^ and Scenic+^13^. These tools are limited by the challenges of deriving accurate gene regulatory networks from incomplete datasets and are restricted to predicting perturbations in transcription factors rather than those in other genes. (2) Learning genetic perturbation embeddings^10, 11, 14, 15^: Methods in this category are designed to map genetic perturbations onto a latent embedding space or incorporate biological priors into deep learning frameworks to elucidate the potential genetic interactions in a specific cell line, thereby enabling the prediction of unseen single or multiple genetic perturbation outcomes; the related approaches include the compositional perturbation autoencoder (CPA)^11, 14, 15^ and the graph-enhanced gene activation and repression simulator (GEARS)^10^. While these methods have demonstrated efficacy in learning genetic perturbation embeddings for prediction purposes, their performance is limited, and their adaptability is constrained when applied to novel and unseen cell lines. (3) Utilizing a single-cell foundation model for genetic perturbation prediction^16-18^: Methods in this category encompass those that can employ large-scale single-cell foundational models via fine-tuning for genetic perturbation prediction. These methods produce generalized and cross-cell-line gene embeddings, suggesting a promising direction for genetic perturbation studies, and these foundation models include scBERT^16^, Geneformer^17^, and scGPT^18^ etc. However, their full potential in the genetic perturbation prediction domain has largely not been explored. Overall, there is still an urgent need to develop a novel strategy that can achieve effective genetic perturbation prediction across various scenarios including single-gene perturbations, multiple-gene perturbations and cross-cell line adaptations, and can accordingly guide proper evaluations of the potential of the existing methods.

It is clear that the three aforementioned test scenarios demand the deployment of innovative strategies to consider several key objectives in genetic perturbation prediction tasks. (1) Modeling high-dimensional genetic perturbation data: The complexity of genetic systems necessitates efficient modeling techniques that can handle the high-dimensional nature of single-cell genetic perturbation data by dissecting and interpreting vast and intricate genetic information. (2) Capturing genetic interactions: This task is critical for effectively capturing potential genetic interactions, either from existing datasets or through the integration of external biological priors^10^. This process is pivotal for understanding the intricate interactions of genes, thereby enabling the prediction of multiple genetic perturbation outcomes. (3) Generalizing to new cell lines: Enhancing the adaptability of computational models to new cell lines is essential. Such generalization is crucial for applying models in diverse biological contexts and ensuring their relevance across varying cellular environments^19^. Overall, the development of a general and robust genetic perturbation model while considering the above three key objectives is challenging.

All these challenges pose great difficulties in terms of directly developing end-to-end models for this task, as previous work have adopted^10, 11, 14, 15^. The fundamental nature of this problem lies in firstly finding genes that can be largely affected by perturbations and their regulatory directions. Then a further goal is to determine the magnitudes of these largely affected genes. Therefore, we realize that this prediction task can be methodically decomposed into these three progressive subtasks in a divide-and-conquer manner. Actually, the concept of decomposing complex problems into subtasks has shown promising results, particularly in scenarios where direct end-to-end deep learning strategies face limitations^20-23^. Notable examples include grounding symbols in visual Sudoku puzzles^24^, unraveling the intricacies of Raven’s progressive matrices^25-27^, and solving mathematical problems using large language models^28, 29^ (LLMs). Previous studies^20,21^ have illuminated the effectiveness of the subtask decomposition approach in tackling such challenges. By partitioning complex problems into a series of smaller, more manageable subtasks and employing a divide-and-conquer strategy, substantial performance improvements have been observed. Furthermore, a positive theoretical result has also shown the benefits of decomposing complex or unlearnable tasks into subtasks in a sequence-to-sequence language model^21^. Taken together, these findings suggest that breaking down a problem into subtasks can significantly enhance the ability of models to learn and solve intricate problems, which inspires us to explore the genetic perturbation prediction problem from the subtask decomposition (STD) perspective.

Herein, we present **<u>S</u>**ub**<u>TA</u>**sk decomposition **<u>M</u>**odeling for genetic **<u>P</u>**erturbation prediction (STAMP), which is a conceptually novel computational strategy inspired by the subtask decomposition learning technique employed by human for complex problems. We formulate the high-dimensional genetic perturbation prediction problem into three progressive subtasks via a divide-and-conquer approach: (1) identifying the postperturbations of differentially expressed genes (DEGs), (2) determining the regulatory directions of these genes, and (3) predicting the magnitudes of the gene expression changes. To this end, STAMP marks a conceptual paradigm shift from the traditional directly end-to-end modeling of the whole task to STD in addressing genetic perturbation prediction, and it is considered as a “plug-in” methodology that can be coupled with any efficient gene embedding approach to solve these subtasks, and the whole perturbation prediction task can be effectively tackled by finding solutions for each of these three subtasks, in line with the inherent logical structure of the problem. This three-subtask structure not only facilitates the design of more targeted predictive models but also serves as a robust and generalizable benchmark for evaluating the performance of various genetic perturbation prediction tools in this domain. Upon benchmarking STAMP against the prevalent methods, STAMP demonstrates substantially superior performance to that of the existing methods in terms of predicting the outcomes of both single and multiple unseen genetic perturbations across distinct datasets, demonstrating the effectiveness of the STAMP strategy. Moreover, STAMP exhibits robust generalizability when applied to previously unseen cell lines across various datasets. Finally, the ability of STAMP to identify key genes and regulatory relationships in new cell lines can be rapidly generalized, and different genetic interaction subtypes can be more precisely detected by enabling more precise and reliable predictions. Overall, STAMP serves as a fundamentally novel and effective prediction and benchmarking strategy that can facilitate genetic perturbation prediction, guide the design of perturbation experiments, and broaden the understanding of perturbation mechanisms.

## Results

### 1. Overview of STAMP

Inspired by the human cognitive strategy of STD used to solve complex problems and its previous successes in complex tasks^20, 21, 29^, we introduce a conceptually novel supervised learning paradigm (Fig. 1a). This paradigm involves partitioning a complex problem into discrete subtasks based on inherent logical structures, where such logical structures clearly exists in single-cell genetic perturbation prediction. Within this context, the problem can be divided into three subtasks following a inherent logical structure: (1) identifying DEG postperturbations, (2) determining the regulatory directions of these genes, and (3) predicting the magnitudes of gene expression changes (Fig. 1a). Therefore, we propose STAMP, an innovative universal framework designed to predict the outcomes of genetic perturbations, including both single genetic perturbations and multiple genetic perturbations within cell line or across cell lines. Central to the functionality of STAMP is its compatibility as a “plug-in” strategy that can be coupled with either a pretrained gene embedding matrix or a dynamically learnable gene embedding matrix, enabling its adaptability to various scenarios. The outputs of STAMP are the outcomes of the three subtasks (Fig. 1b).

**Fig. 1:**
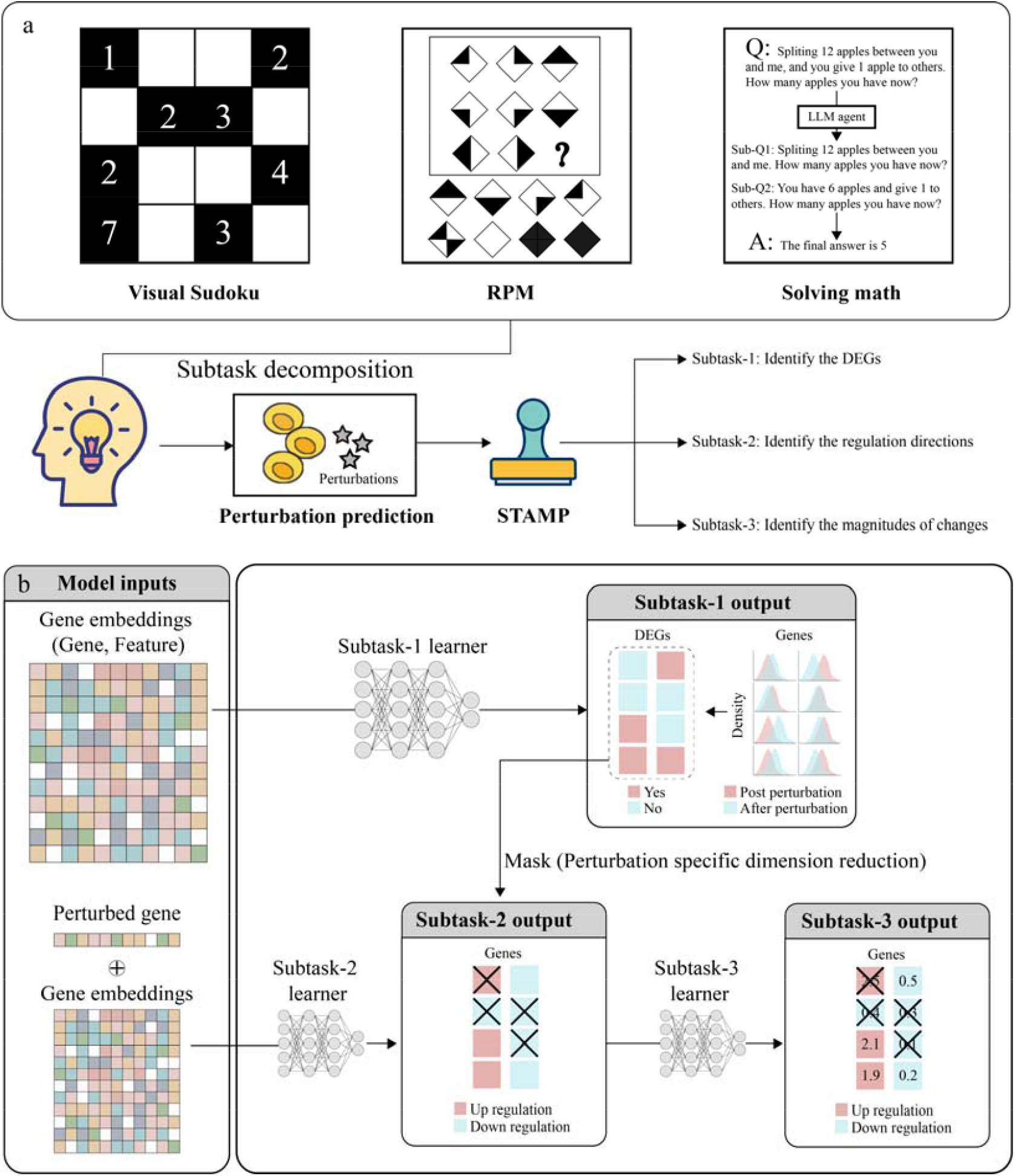
Illustration of the STAMP framework. a. Inspired by the STD technique used by humans to solve for complex problems and the pioneering success achieved in tasks such as solving the visual sudoku problem, unraveling Raven’s progressive matrices and solving math problems with LLMs, the genetic perturbation prediction task can be decomposed into three subtasks by our proposed STAMP framework. b. STAMP model architecture. STAMP takes gene embeddings as its inputs, and the learner of subtask-1 outputs DEG postperturbations. The learner of subtask-2 outputs the regulatory directions of the DEG postperturbations. The learner of subtask-3 takes the output of subtask-2 as an intermediate supervisory signal and outputs the magnitudes of the DEG changes.

Specifically, STAMP is constructed on the basis of gene embeddings that capture the intrinsic biological nature of genetic interactions (Fig. 1b). STAMP strategically divides this perturbation prediction problem into three subtasks, each of which leverages these gene embeddings to unravel different aspects of genetic perturbations. In the first subtask, STAMP utilizes the gene embeddings of perturbed genes to predict DEG postperturbations. This process effectively constructs a mapping from the gene embedding space to the corresponding DEG postperturbation space (Fig. 1b). Given that the majority of genes remain unaffected by perturbations in most cases, the changes of gene expression vectors postperturbations characteristically exhibit high-dimensional sparsity. Therefore, the DEGs identified in the first subtask can be considered as a perturbation-specific dimension reduction process, helping to improve the signal-to-noise ratio in data. The second subtask further exploits this special property and employs the gene embeddings to predict the regulatory directions of the DEG postperturbations (Fig. 1b). In this subtask, STAMP extends the mapping from the gene embedding space to a new dimension, focusing on the regulatory direction of the postperturbation space. This step is pivotal for delineating the regulatory trajectories of DEGs, whether they are upregulated or downregulated following perturbation. The third subtask involves the use of gene embeddings to predict the magnitudes of gene expression changes (Fig. 1b). This subtask aids in quantifying the extents of expression alterations, providing a quantitative picture of the impacts of genetic perturbations. During the model optimization process, STAMP harnesses the power of multitask supervised learning^30^ in combination with intermediate supervised learning techniques to jointly solve the three subtasks via gradient descent by considering two key optimization steps: 1) jointly learning the first and second subtasks; and 2) setting the second subtask as the intermediate supervisory signal for the third subtask. To this end, this three-subtask structure not only enhances the prediction accuracy of the model but also establishes a more rigorous and generalizable evaluation system for assessing methodologies within the domain of genetic perturbation prediction.

### 2. Challenge 1: Predicting single genetic perturbation outcomes

For evaluating the proposed approach in the context of single genetic perturbations, a model was constructed on the basis of gene perturbations that were excluded from the training dataset, ensuring that the predictive capabilities of this model were tested on previously unseen genes. This comparison utilized preprocessed data from four distinct datasets: the RPE1_essential dataset^31^ with 1694 perturbations, the K562_essential dataset^31^ with 1741 perturbations, the TFAtlas dataset^32^ with 1183 perturbations and the K562_GW dataset^31^ with 7131 perturbations (Supplementary Notes 1-4, Supplementary Table 1). Each model was trained separately on each dataset via fivefold cross-validation (Supplementary Note 6). STAMP was designed to adapt both well-pretrained gene embeddings and learnable gene embeddings. Specifically, we utilized prominent well-pretrained gene embeddings from foundational models such as scBERT, Geneformer and scGPT (these models are denoted as “scBERT+STAMP”, “Geneformer+STAMP”, and “scGPT+STAMP”, respectively) and dynamic learnable gene embeddings in the GEARS with a network structure (this model is denoted as “GEARS+STAMP”) as the inputs of STAMP (Supplementary Note 5). For the GEARS, we extended the model by integrating STAMP into its outcome prediction component. This integration step was designed to leverage the advanced predictive abilities of STAMP within the existing GEARS framework (Supplementary Note 5). Furthermore, to establish a comprehensive and rigorous evaluation, we included random gene embeddings (this model is denoted as “Random+STAMP”), the original GEARS (this model is denoted as “GEARS”), and the CPA as baseline models for comparison purposes. Notably, the CPA was originally designed for predicting the outcomes of compositional perturbations, of which single perturbations were observed during training. We modified this model by substituting its learnable gene embedding component with various well-pretrained gene embeddings (these models denoted are as “scBERT+CPA”, “Geneformer+CPA”, and “scGPT+CPA”; see Supplementary Note 5). This adjustment enabled the CPA to predict outcomes for previously unseen perturbations, thus facilitating fair comparisons.

The ground truths of the DEGs, their regulatory directions and the fold changes observed after perturbations in the three subtasks were identified by a standard statistical analysis^33, 34^ conducted on each dataset (Fig. 2a, Supplementary Note 4). As the identification of DEG postperturbations in the first subtask and the determination of the regulatory directions of these DEGs in the second subtask are both binary classification tasks, we evaluated the performance of the rested models by calculating the area under the receiver operating characteristic curve (ROC-AUC) and the area under the precision-recall curve (PR-AUC) for both the first subtask and the second subtask. In the third subtask, we measured the Pearson correlation coefficient^35^ (PCC) and mean squared error (MSE)^36^. Furthermore, considering that the majority of genes exhibit small variations between their unperturbed and perturbed states, our evaluations in the second and third subtasks were further confined to more challenging settings. Specifically, we focused on the evaluation of the top 20 most differentially upregulated genes and the top 20 most differentially downregulated genes (top 40 DEGs; Supplementary Note 4). Overall, the test results indicate that when coupled with the STAMP strategy, models such as the GEARS or gene embeddings from large foundation models showed remarkable improvements across the three subtasks and testing scenarios involving the top 40 DEGs (Figs. 2b-d, Extended Data Fig. 2, Supplementary Note 7, Supplementary Tables 3-10, Supplementary Figs. 1-3). Additionally, the poor performance of the CPA with pretrained gene embeddings (“scBERT+CPA “, “Geneformer+CPA” and “scGPT+CPA”) indicated that the STAMP strategy is necessary for achieving improved genetic perturbation prediction. Moreover, benchmark tests conducted on different gene embeddings within perturbation prediction tasks revealed that the “GEARS+STAMP” and “scGPT+STAMP” models consistently outperformed other large foundational models across all testing scenarios. In addition, when comparing the time costs of the methods, “scGPT+STAMP” had a faster computation speed than “GEARS+STAMP” (Supplementary Table 2). Collectively, these findings highlight that both well-defined gene embeddings and the implementation of the STAMP strategy are essential prerequisites for accurately predicting single genetic perturbation outcomes.

**Fig. 2:**
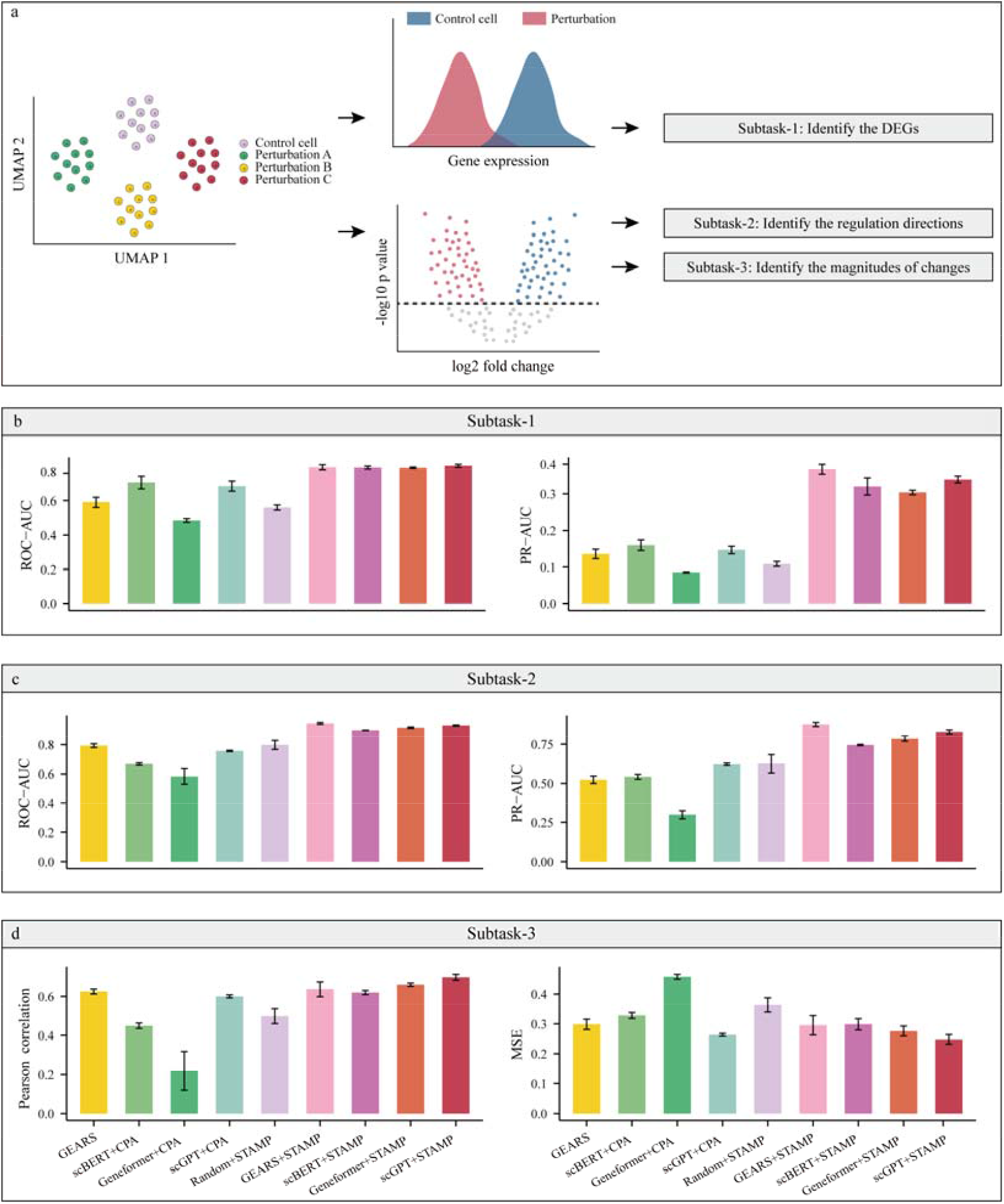
Challenge 1. Predicting single genetic perturbation outcomes in the RPE1_essential dataset. a. The ground truths of three subtasks were identified through a standard single-cell statistical analysis. b. The ROC-AUC and PR-AUC values of different models in subtask-1. c. The ROC-AUC and PR-AUC values of different models in subtask-2. e. The Pearson correlations and MSEs of different models in subtask-3. The data are presented as their mean values, and the error bars show the 95% confidence intervals (CIs) of the mean estimates.

### 3. Challenge 2: Predicting multiple genetic perturbation outcomes

STAMP has a remarkable ability to predict the outcomes of multiple genetic perturbations. This is achieved by transforming the embeddings of multiple genes into inputs for the predictive model. However, it should be noted that in the available datasets for multiple genetic perturbation scenarios, two-gene perturbations are the predominant components. Therefore, to address the challenge of predicting multiple genetic perturbations, the model evaluation was centered around two-gene perturbation scenarios. This prediction scenario could be categorized into three distinct cases, each defined by the extent of the model’s exposure to the genes during training (Fig. 3a). The first case encompassed instances where the model had previously encountered each of the two genes within the training data (zero of two unseen). In the second case, the model was exposed to only one of the two genes in the perturbation pair during the training phase (one of two unseen). The third case was the most challenging scenario, where neither of the two genes in the combination were presented in the training data (two of two unseen). To conduct this comprehensive comparison, we utilized preprocessed data from three distinct datasets: the PRJNA551220 dataset^5^ with 226 perturbations, the Perturb-CITE-seq dataset^37^ with 1360 perturbations and the PRJNA787633 dataset^38^ with 1573 perturbations. Each model was trained separately on each dataset via fivefold cross-validation (Supplementary Note 1, Supplementary Table 1).

**Fig. 3:**
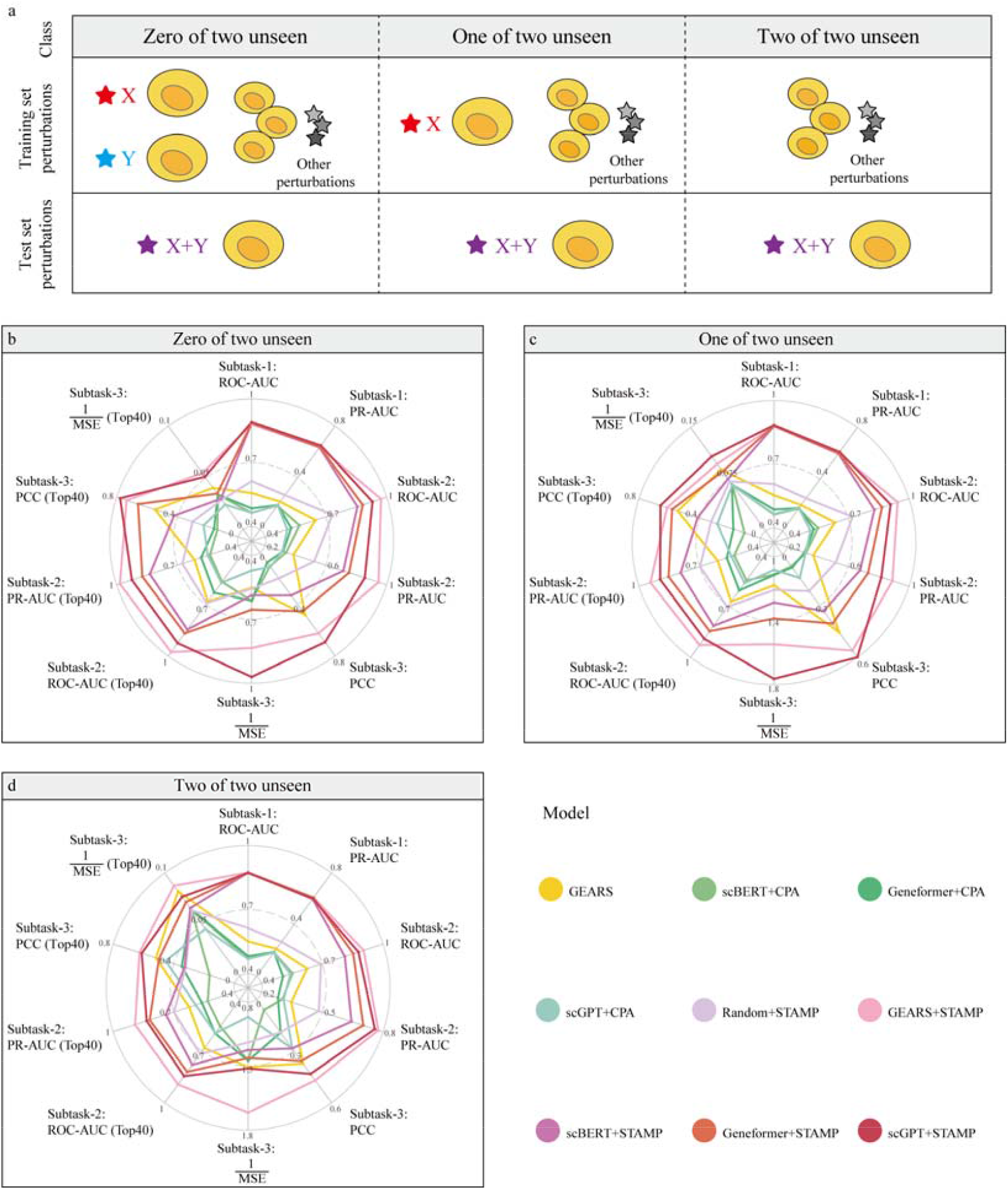
Challenge 2. Predicting multiple genetic perturbation outcomes in the PRJNA551220 dataset. a. Illustration of the three testing scenarios for evaluating the two-gene perturbation prediction capabilities of models. X and Y represent different single genetic perturbations. X+Y represents the multiple genetic perturbations caused by perturbations X and Y. b. Performance of different models in terms of different performance metrics, including metrics considering the top 40 DEGs, in the scenario with zero of two perturbations unseen. c. Performance of different models in terms of different performance metrics, including metrics considering the top 40 DEGs, in the scenario with one of two perturbations unseen. d. Performance of different models in terms of different performance metrics, including metrics considering the top 40 DEGs, in the scenario with two of two perturbations unseen. The data are presented as their mean values.

For each dataset, the ground truths of the three subtasks were carefully determined by conducting a standard statistical analysis for each specific prediction case (Supplementary Note 4). Subsequently, each model underwent a comprehensive evaluation across the three subtasks for every prediction case. Additionally, a focused evaluation was conducted on the second and third subtasks, particularly for the top 40 DEGs identified in each prediction case. This multilevel assessment strategy ensured a thorough and nuanced evaluation of the predictive capabilities of the models. As a result, consistent with the findings of challenge 1, the models or gene embeddings derived from large foundational models and coupled with the STAMP strategy exhibited substantial improvements across all testing scenarios (Figs. 3b-d, Supplementary information). Among these, the “GEARS+STAMP” and “scGPT+STAMP” models displayed the best performance, yielding better performance than those of the other foundational models (Supplementary Tables 11-28, Supplementary Figs. 4-5). In addition, when comparing the time costs of the models, “scGPT+STAMP” had a faster computational speed than “GEARS+STAMP” (Supplementary Table 2). Collectively, these findings again highlight that coupling a model with the STAMP strategy is an essential prerequisite for accurately predicting multiple genetic perturbation outcomes.

### 4. Challenge 3: Predicting genetic perturbation outcomes across cell lines

A significant challenge in the domain of genetic perturbation prediction across cell lines is the variability in the genetic interaction patterns across different cell lines (Fig. 4a). This heterogeneity often results in models trained on one cell line dataset exhibiting poor generalizability to other cell line datasets. Few-shot learning and zero-shot learning represent prominent methods in domain adaptation areas^19, 39, 40^. However, such interaction pattern diversity in different cell lines complicates the feasibility of performing zero-shot prediction^39^ across cell lines. Further compounding this issue is the limited availability of diverse cell line datasets. This scarcity hinders the effective application of advanced techniques such as meta-learning^40, 41^, which are currently infeasible due to dataset constraints. Therefore, the key question here is how quickly a predictive model can adapt to a novel cell line. To address this difficult challenge, we conducted an in-depth evaluation of the ability of STAMP to predict the outcomes of genetic perturbations in a novel cell line dataset for which very few samples (few-shot setting), including zero samples (zero-shot setting), were available for training (Supplementary Note 9). In this evaluation, we utilized the data from 8 distinct datasets and preprocessed data, including 3 CRISPRi datasets and 5. CRISPRa datasets. The 3 CRISPRi datasets consisted of the K562_GW dataset^31^, RPE1_essential dataset^31^ and Perturb-CITE-seq dataset^37^. The 5 CRISPRa datasets consisted of the TFatlas dataset, the PRJNA551220 dataset^5^, the PRJNA787633 dataset^38^, the PRJNA641125a dataset^42^ and the PRJNA609688 dataset^43^ (Supplementary Note 8).

**Fig. 4:**
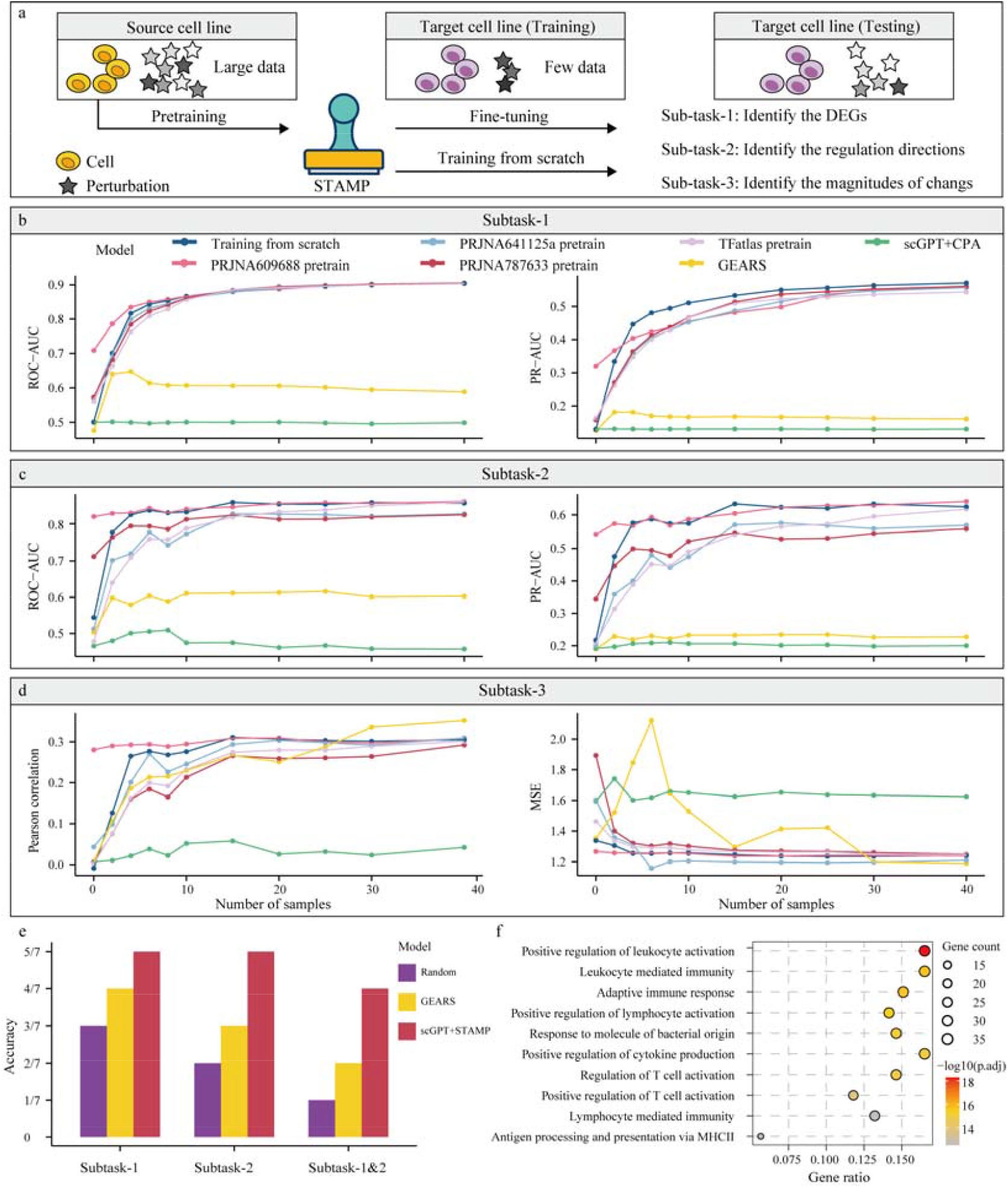
Challenge 3. Predicting genetic perturbation outcomes across cell lines. a. Illustration of the testing scenario implemented across different cell line datasets. When generalizing to a new cell line dataset, STAMP can use the pretrained parameters from an source cell line dataset and then be fine-tuned with few samples from the target cell line dataset. STAMP can also be trained from scratch with few samples from the target cell line dataset. b. The ROC-AUC and PR-AUC values produced by different models on pretrained cell line datasets in subtask-1. c. The ROC-AUC and PR-AUC values produced by different models on pretrained cell line datasets in subtask-2. d. The Pearson correlations and MSEs yielded by different models on pretrained cell line datasets in subtask-3. A performance comparison was conducted on the PRJNA551220 dataset. The name of dataset represents the “scGPT+STAMP” pretrained on that dataset (for example, “PRJNA609688 pretrain” denotes the “scGPT+STAMP” model pretrained on the PRJNA609688 dataset.) Each model was fine-tuned with various numbers of samples from the target datasets (0, 2, 4, 6, 8, 10, 15, 20, 25, 30, and 40 samples). The plot shows the average performance as a function of the number of samples utilized from the target cell line dataset. e. The accuracies achieved by different models in three testing scenarios, i.e., the first subtask, the second subtask, and the testing scenario where both the first subtask and the second subtask needed to be correct. f. GO analysis of the DEGs predicted by “scGPT+STAMP” after perturbing the IFNGR1 gene.

For each novel cell line, two test scenarios are applicable, i.e. few-shot learning (few perturbation samples available) and zero-shot learning (no perturbation samples available). For few-shot learning, we adopted two distinct strategies to evaluate the performance of STAMP: (1) Training STAMP from scratch with random initialization, using a small number of available perturbations for the novel cell lines. Evaluating the first strategy is the most straightforward way to ascertain the ability of a model to learn and predict perturbation outcomes from a constrained dataset, which is a common scenario in practical applications. (2) Fine-tuning STAMP, initially pretrained on a dataset from a different cell line, on the same small set of perturbations available for the novel cell lines. For zero-shot learning, only the second strategy is applicable while the zero-shot prediction is conducted by directly predicting outcomes for novel cell lines using pretrained STAMP without the need for any finetuning. Noted that the evaluations of the second strategy in two test scenarios were constrained by the use of cell lines with the same CRISPR type. During this process, one of the cell line datasets (the source dataset) was used to pretrain the model and generalize it to predict the novel cell line datasets (the target datasets). To this end, STAMP was trained in two phases: on all perturbations from the source cell line dataset (pretraining phase) and then on the small number of perturbations available from one of the target cell line datasets (fine-tuning phase) or zero number of perturbations available for zero-shot learning without fine-tuning. Note that in all the tests involving the second strategy, for simplicity, we use the name of the dataset to represent the model (for example, “PRJNA609688 pretrain” denotes the STAMP model pretrained on the PRJNA609688 dataset and fine-tuned with a small set of samples from the target dataset). As “scGPT+STAMP” showed superior performance and fast computational speeds in our previous study, we used it as the representative STAMP model in this challenging evaluation (Supplementary Table 2). Furthermore, to establish a comprehensive comparison, we included “GEARS” and “scGPT+CPA” as baseline models for comparison purposes. Each evaluation was conducted via fivefold cross-validation, and the model evaluation was conducted across the three subtasks. Moreover, a focused evaluation was also conducted on the second and third subtasks, particularly for the top 40 DEGs.

In the evaluations of the first strategy, the performance of “GEARS” and “scGPT+CPA” improved slowly as samples from the new cell line dataset were added to the training set. In contrast, “scGPT+STAMP” improved rapidly, with a large increase in performance after examining only an additional 10-20 samples in most cases (Figs. 4b-d, Extended Data Fig. 4, Supplementary Figs. 6-12). In the evaluations of the second strategy, we evaluated the performance of “scGPT+STAMP” when it was pretrained on different source datasets and subsequently applied to a target dataset (Figs. 4b-d, Extended Data Fig. 4, Supplementary Figs. 6-12). Interestingly, although the pretrained “scGPT+STAMP” model sometimes exhibited superior performance in the zero-shot setting^39^ (sample number =0) compared to that of training “scGPT+STAMP” from scratch, the latter often outperformed the former in the few-shot setting^44^ (sample number > 0; Supplementary Figs. 6-12). This discrepancy could be attributed to the gene interaction pattern differences across different cell lines. The presence of consistent gene interaction patterns between certain cell lines contributed to the enhanced performance of the pretrained STAMP model in the zero-shot setting, surpassing that of the randomly initialized models. However, differences in the interaction patterns of certain gene pairs could lead to a nonbenign parameter initialization process, limiting the improvement exhibited by the model in the few-shot setting. Notably, this phenomenon diminished when the model was both trained and evaluated within the same cell line (Figs. 4b-d, Extended Data Fig. 4), where the PRJNA609688 dataset and the PRJNA551220 dataset represented the same cell line from distinct sequencing experiments. When conducting the evaluations on the PRJNA609688 dataset and the PRJNA551220 dataset, STAMP was pretrained on one of these two datasets, and it consistently achieved robust performance in both zero-shot and few-shot settings (Figs. 4b-d, Extended Data Fig. 4). Further exploration into the correlation between the zero-shot performance of pretrained models and cell line similarity revealed a notable trend: models pretrained on cell line datasets with higher degrees of similarity tend to exhibit enhanced performance in the zero-shot setting (Supplementary Fig. 13). Collectively, these findings highlight the ability of STAMP to adapt across cell lines and its robustness to experimental batch effects^45^.

### 5. STAMP uncovers key regulatory genes and pathways with small samples

To further demonstrate that STAMP can quickly adapt to a novel cell line, we applied “scGPT+STAMP” to the scCRISPR-seq dataset^46^, for which only 24 perturbations were available (Supplementary Note 8). On this dataset^46^, Papalexi et al. applied ECCITE-seq^47^ to explore the molecular networks that regulate PD-L1 expression and reported that IFNGR1, IFNGR2, IRF1, JAK2, and STAT1 are positive regulators of PD-L1, while BRD4 and CUL3 are negative regulators of PD-L1^46^ (Supplementary Note 8). Therefore, in this scenario we examined whether STAMP could accurately predict the effects of the 7 perturbations on PD-L1 expressions with a small set of perturbations. As the first strategy (Training STAMP from scratch) exhibited superior performance in few-shot setting and it is a more straightforward way in practical scenarios, it was adopted in this dataset. STAMP without training (denoted as “Random” in this scenario) and “GEARS” were used as baseline models for comparison purposes. As the Pearson correlation coefficient and MSE cannot be calculated with only a single gene expression value, the performances of the baseline “scGPT+STAMP”, “GEARS”, and “Random” models were examined only in the first and second subtasks. In the first subtask, if the predicted score of PD-L1 for a certain gene perturbation was above the 80th percentile, then PD-L1 was classified as a differentially expressed gene for that perturbation (using the 85th, 90th, 95th percentiles, we arrived at similar conclusions; Supplementary Fig. 13). In the second subtask, the 50th percentile was used as the threshold for defining the direction of regulation.

As the results showed, in the first subtask, “scGPT+STAMP” successfully predicted that 5 out of these 7 genetic perturbations had significant effects on PD-L1 expression, outperforming the “GEARS” and “Random” baseline models (Fig. 4e). Furthermore, “scGPT+STAMP” achieved the best performance in the second subtask (Fig. 4e). In addition, we evaluated the performance of STAMP in a more challenging scenario where both the first and second subtasks for perturbation prediction needed to be correct. “scGPT+STAMP” successfully predicted 4 (IFNGR1, IFNGR2, IRF1, and JAK2) out of the 7 perturbations, which was twice the success rate of “GEARS” (Fig. 4e). In subsequent analyses, we explored the mechanisms underlying the downregulation of PD-L1 expression. For each of these 4 successfully predicted genetic perturbations, all DEGs in the first subtask identified by “scGPT+STAMP” were subjected to a gene ontology (GO) analysis^48^ (Supplementary Note 10). Our findings revealed that these perturbations led to the inhibition of immune-related pathways (Fig. 4f and Supplementary Fig. 14), resulting in a decrease in PD-L1 expression. This outcome aligned with the findings from prior research^46, 49, 50^, further validating the predictive accuracy of our model. In summary, the above results demonstrate that STAMP can quickly adapt to a novel cell line, which is essential in the perturbation prediction field because perturbation sequencing technology is still in its early stage and cost-ineffective, and many cell lines have limited perturbation data.

### 6. STAMP precisely uncovers diverse types of genetic interactions

In scenarios involving two-gene perturbations, a simple way to predict the outcome of a combinatorial perturbation might involve summing the individual effects of perturbing each gene separately. However, this approach is primarily applicable when the gene interactions (GI) are additive, where they are referred to as additive interactions^5^ (Supplementary Note 11). Biological reality is often complex, with genes known to engage in various forms of nonadditive interactions following perturbation. These interactions, which have been defined in previous studies^5^, include subtypes such as synergy, suppression, neomorphism, redundancy and epistasis^10^ (Fig. 5b, Supplementary Note 12). Genetic interaction scores (GI scores), such as magnitudes, model fitting scores, the equality of contributions and similarity scores, have been defined to identify these different genetic interaction subtypes^5, 10^ (Supplementary information). Previous work^10^ has advocated for the use of a single GI score threshold to categorize different GI interaction subtypes (Supplementary Note 11). In this study, we evaluated the performance of “scGPT+STAMP” and “GEARS” in terms of identifying new genetic interactions. To estimate the upper performance limit, we also calculated the GI scores based on the ground-truth gene expression values^10^ (Ground truth in Fig.5) and collected the GI scores calculated in the original study^5^ for all combinatorial genes (Ori_study in Fig.5, Supplementary Note 12). We also utilized Precision@10 to evaluate the performance of the models (Supplementary Note 13). This metric calculates the proportion of predicted combinations within the top ten that truly exhibit a specific genetic interaction subtype, as reported by the original study. Compared to “GEARS”, “scGPT+STAMP” improved the Precision@10 metrics obtained for four out of five GI subtypes, and the improvements exceeded 50% for neomorphism, redundancy and epistasis (Fig. 5a).

**Fig. 5:**
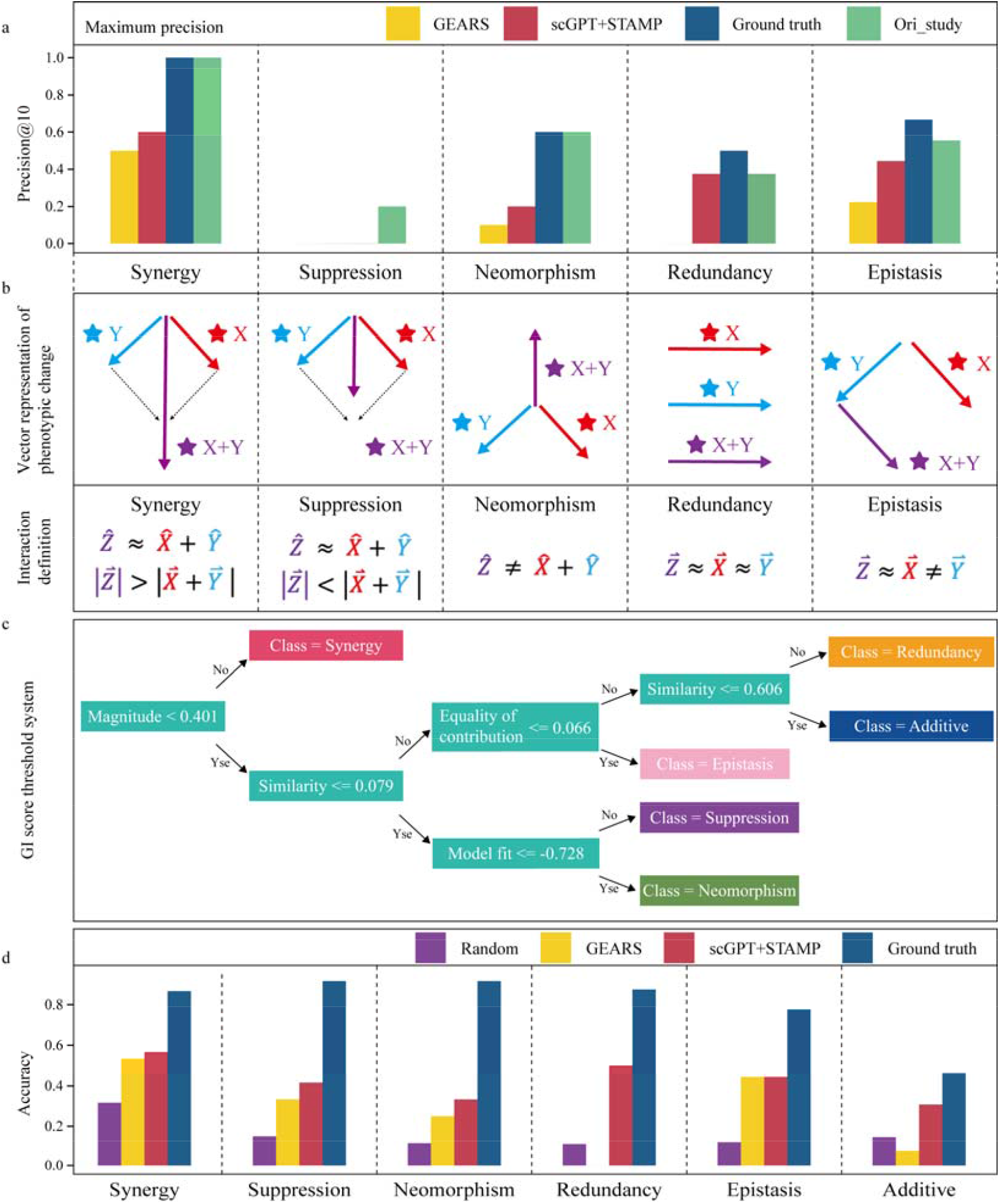
Genetic interaction subtype identification results of STAMP. a. Precision@10 scores obtained when predicting genetic interactions from 125 two-gene perturbations with GI scores obtained by STAMP (“scGPT+STAMP”), GEARS (“GEARS”), the ground-truth gene expression vector (“Ground truth”) and the original study (“Ori_study”). b. Definitions of genetic interaction subtypes. c. The decision tree of the GI score threshold system was constructed from the ground-truth gene expression vector, and each GI score was normalized to a z score. d. Comparison among the accuracies of different models in terms of predicting genetic interactions with the decision tree constructed in c.

However, all GI scores from different models and sources showed that the tested models struggled to identify suppression and redundancy subtypes (Fig. 5a). Notably, only the GI scores from the original study had a Precision@10 of 0.2 for suppression, and the GI scores from the original study had a Precision@10 of 0.4 for redundancy (Fig. 5a). This indicated that relying on a single GI score is insufficient for interpreting all GI interaction subtypes and highlighted the importance of designing a more suitable GI score threshold system to accurately identify GI subtypes, as a single GI score threshold falls short of capturing the complexity inherent in these GIs (Fig. 5a, Supplementary Notes 12-13).

To develop a more accurate GI score threshold system for identifying GI subtypes, we constructed a decision tree^51^ model utilizing GI scores calculated from ground-truth gene expression values (Supplementary Notes 12-13). This approach was chosen due to the superior explanatory power of decision trees, facilitating a clearer understanding of the classification criteria and underlying patterns in the data (Fig. 5c). Based on the decision tree model, we observed that the upper accuracy limit of identifying different GI subtypes based on the ground-truth gene expression values exceeded 0.75 across all nonadditive GI subtypes. Notably, this accuracy exceeded 0.85 for four out of the five nonadditive subtypes (Fig. 5d). When “scGPT+STAMP” was compared with “GEARS” and the “Random” baseline, we also incorporated additive interactions into this evaluation system. STAMP outperformed the other models for five out of the six GI subtypes, while STAMP exhibited comparable accuracy to that of “GEARS” for the epistasis subtype (Fig. 5d). Collectively, our findings indicate that the newly designed GI score threshold system offers a more accurate evaluation of the effectiveness of different models in terms of identifying GI subtypes and that STAMP exhibits a superior ability to accurately discern these GIs, underscoring its potential role as a robust model in this domain.

## Discussion

The rapid development of single-cell transcriptome-based high-throughput sequencing and CRISPR-based perturbation techniques has enabled the examination of perturbation effects after target genes are perturbed. However, due to the cost of sequencing, the vast combinatorial search space and the heterogeneity of the genetic interaction patterns within various cell lines, the available data are limited compared to the full sample space. This highlights the need for computational methods that can accurately predict perturbation outcomes across various scenarios such as those involving single genetic perturbations, multiple genetic perturbations and perturbations across cell lines. Therefore, we introduce STAMP, a conceptual paradigm shift from the traditional directly end-to-end modeling of the whole task to STD in addressing genetic perturbation prediction. The genetic perturbation prediction task can naturally be decomposed into three subtasks following the inherent logical structure under the whole task: identifying DEG postperturbations, determining the regulatory directions of DEGs, and predicting the magnitudes of gene expression changes. STAMP not only is considered as a “plug-in” methodology that can be coupled with any efficient gene embedding to start solving these subtasks but also serves as a robust and generalizable benchmark framework for evaluating the prediction performance of models constructed for the perturbation prediction task. Accurately predicting the outcomes of genetic perturbations involves three challenges: (1) predicting single genetic perturbation outcomes; (2) predicting multiple genetic perturbation outcomes; and (3) predicting genetic perturbation outcomes across cell lines. We examined models for addressing all three challenges through our three-subtask benchmark framework, and STAMP exhibited superior performance to that of the other methods. Furthermore, the ability of STAMP to identify key genes and regulatory relationships in new cell lines can be rapidly generalized, and STAMP can also serve as a robust tool for precisely identifying genetic interaction subtypes.

Nevertheless, the efficacy of STAMP is currently constrained by the availability of data. To obtain reliable predictions, particularly when generalizing to new cell lines, STAMP requires a few samples for training or fine-tuning the model (few-shot setting). Although we examined the zero-shot setting for entirely new cell lines, STAMP still has a great improvement space in this case. Future iterations of STAMP are anticipated to overcome these limitations as more comprehensive perturbation prediction data encompassing a diverse array of cell lines become available. This data expansion will be crucial for enhancing the predictive capacity of STAMP and broadening its applicability in the field of genetic research. Furthermore, the development of a carefully designed output schema for the STAMP model holds significant potential. By designing the model to output a confidence interval for each subtask, it becomes possible to capture and quantify the distributional changes in each gene. Moreover, the presence of “escaping” cells^46, 52^ that evade the effects of a perturbation may influence the estimation of perturbation effects. Future work will identify ground truths for the three subtasks via various DEG identification pipelines while accounting for these “escaping” cells; this could further improve the performance of STAMP.

Notably, our findings underscore the critical role of gene embeddings in accurately predicting genetic perturbation outcomes. As the existing foundational models are pretrained on a large repository of normal human cells, the gene interaction patterns among diseased cells may not be effectively captured. A well-defined gene embedding method that is capable of comprehensively capturing the potential relationships among genes can significantly enhance the performance of STAMP. Consequently, the pursuit of developing refined gene embeddings that are both specific to individual cell lines and applicable across various cell lines remains an essential avenue for further research.

In conclusion, STAMP represents a significant advancement in the field of genetic perturbation prediction and it underscores the effectiveness of the subtask decomposition-based learning as a novel AI paradigm in biological domain. Specifically, its methodical approach to decomposing the complex genetic perturbation prediction task into three subtasks provides a blueprint for future research in this task. Future work involving the application of STAMP to the drug perturbation prediction task is anticipated. As the field of bioinformatics continues to evolve, tools such as STAMP will be instrumental in unlocking the intricacies of genetic networks and their therapeutic implications for human health.

## Methods

### 1. Overview of STAMP

STAMP considers a perturbation dataset comprising *N* cells, denoted as 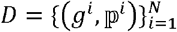, where *g*^*i*^ ∈ *R*^*K*^ represents the gene expression vector of cell *i*, which encompasses *K* genes. Furthermore, 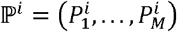 constitutes the set of *M*-sized perturbations performed on cell *i*. Each perturbation 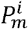 within this set correlates with the index of a specific gene. Given a set of perturbations ℙ ^*i*^, the primary objective of STAMP is to predict the outcomes of perturbations through three distinct subtasks, including identifying the DEGs postperturbations, determining the regulatory directions of these DEGs and quantifying the magnitudes of the gene expression changes exhibited by these DEGs.

STAMP is constructed on the gene embeddings, denoted as *Z*, which also serve as the inputs of the model. To better capture the potential genetic interactions for gene embedding, we represent each gene *u*_*i*_ as an embedding *z*_*i*_ ∈ *R*^*d*^ via a gene encoder *f*_*gene*_ : *u* → *R*^*d*^ Because each gene has its own perturbation pattern, in the first subtask, STAMP employs a gene response predictor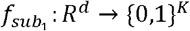, which aims to learn the mapping from the *d*-dimensional gene embedding space to a *K*-dimensional postperturbation DEG space via a series of perturbation sets ℙ. Subsequently, for the second subtask, STAMP utilizes a gene regulatory predictor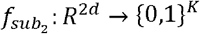. This predictor aims to learn the mapping from the 2*d*-dimensional combined gene embedding space to a *K*-dimensional postperturbation regulatory direction space via a series of perturbation sets ℙ. Moreover, the predictor in the second subtask is further leveraged as intermediate supervisory information to aid in constructing the gene expression change predictor 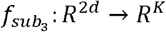. This mapping translates gene embeddings to vectors that are indicative of postperturbation gene expression changes.

### 2. Gene response predictor in subtask-1

To accurately predict outcomes for genes perturbed in a previously unseen scenario, STAMP requires gene embeddings that effectively capture genetic interactions. Specifically, STAMP can utilize either well-established gene embeddings derived from large single-cell foundational models such as the scBERT, GeneFormer and scGPT or learnable gene embeddings from gene encoders, such as the GEARS. Given a perturbation set ℙ =(*P*_1_,*P*_2_,…*P*_*M*_), STAMP retrieves the gene embedding of each element of that set, resulting in (*Z*_1_,*Z*_2_,…*Z*_*M*_). To model multiple genetic perturbations, the ‘mean’ operator is employed to represent the combined representation of multiple genes. This operator provides flexibility and extendibility to perturbations of various sizes. Consequently, the perturbation embedding for this set, denoted by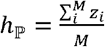, is then input into the gene response predictor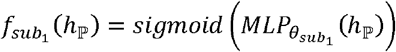 for the first subtask.

Given that most genes do not exhibit significant changes in their postperturbation expression levels, the class labels in the first subtask, which denote whether they are postperturbation DEGs, are inherently imbalanced. To address this issue, we implement a batch-level reweighting loss that assigns greater weights to the DEGs. For a minibatch consisting of *T* perturbations, each perturbation ℙ_*k*_ has *T*_*k*_ cells, and each cell has *K* genes. The ground-truth DEGs 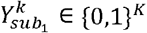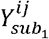 are identified by standard statistical analysis techniques that are commonly used in single-cell studies. The loss for this subtask is defined as follows:

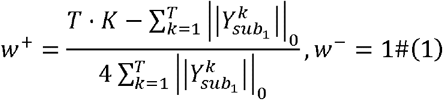

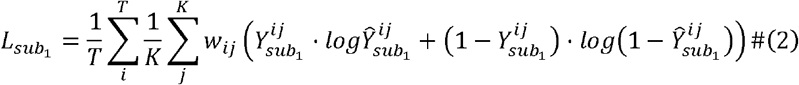

where *w* ^*+*^ is the weight of the positive class and *w* ^*−*^ is the weight of the negative class. *w*_*ij*_ denotes the weight of gene *j* in batch *i*, denotes the ground-truth label of gene *j* in batch *i* and 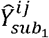 denotes the predicted class score of gene *j* in batch *i*.

### 3. Gene regulatory predictor in subtask-2

To accurately capture the intricate regulatory directions between perturbed genes and other genes, a perturbation embedding *h*_ℙ_ is applied to every gene embedding *z*_*i*_ to obtain a perturbation-induced, combined gene embedding. For gene *u*, this results in a combined embedding *h*_ℙ_║*z*_*u*_, where ║ denotes the concatenation operation. Subsequently, this combined embedding for each gene is input into a learnable multilayer perceptron (MLP), generating a postperturbation gene embedding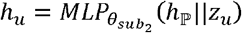. In subtask-2, postperturbation gene embedding is subsequently input into a gene-specific layer, which is parameterized by 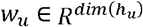, *b*_*u*_, to map each gene to a scalar value representing the corresponding regulatory direction. The regulatory direction for each gene is thus determined as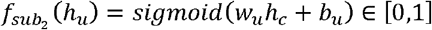. Different datasets may introduce varying degrees of imbalance in the class labels for the second subtask, where these labels denote whether genes are upregulated or downregulated after experiencing perturbations. To address this, we also implement a batch-level reweighting loss that assigns greater weights to genes that are less represented in the dataset. Notably, we focus on the regulatory directions of the genes among the DEGs from the first subtask. A minibatch consists of *T* perturbations, where each perturbation ℙ_*k*_ has *T*_*k*_ cells and each cell has *K* genes. The ground-truth regulatory directions of the genes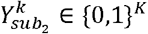 are identified by the standard statistical analysis techniques that are commonly used in single-cell studies. The loss for this subtask is defined as follows:

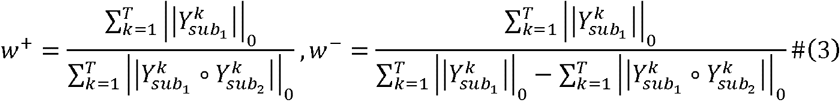

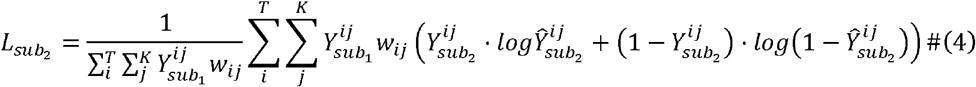

where *w*^*+*^ is the weight of the positive class and *w*^−^ is the weight of the negative class. *w*_*ij*_ denotes the weight of gene *j* in batch *i*.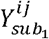 denotes whether gene *j* in batch *i* is a DEG, and 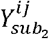 denotes whether gene *j* in batch *i* is upregulated or downregulated.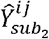 denotes the predicted class score of gene *j* in batch *i*.

### 4. Gene expression change predictor in subtask-3

Relative to the more nuanced magnitudes of gene expression changes, the directions of gene regulation represent coarser outcomes, essentially capturing the binary positive or negative natures of gene expression changes. In subtask-3 of STAMP, we address this issue by processing the postperturbation gene embedding 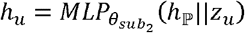 for each gene *u* through a feature extraction layer. This layer is designed to discern the intricate and subtle features that are embedded within postperturbation gene expression profiles. The refined postperturbation gene embedding, denoted as 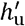, is obtained through 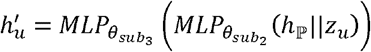. For each gene *u*, the associated refined postperturbation gene embedding is then input into a gene-specific layer, which is parameterized by 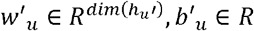, to map each gene to a scalar value representing the magnitude of the gene expression change. As we use the fold change value to quantify the gene expression change, the change for each gene is thus determined as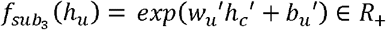. For a minibatch consisting of *T* perturbations, each perturbation ℙ_*k*_ has *T*_*k*_ cells, and each cell has *K* genes. The ground-truth gene expression changes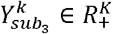 are identified by the mean fold change calculation based on the unperturbed cells. The loss for this subtask is defined as follows:

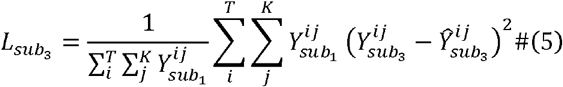

where 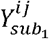 denotes whether gene *j* in batch *i* is a DEG and 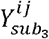 denotes the mean fold change value of gene *j* in batch *i* .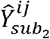 denotes the predicted fold change value of gene *j* in batch *i*.

Then, the overall prediction loss function is 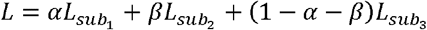, and we take *α* = 0.1 and *β* = 0.5 in this study (Supplementary Note 14).

## Acknowledgements

This work was supported by the National Key Research and Development Program of China (Grant No. 2021YFF1201200, No. 2021YFF1200900), National Natural Science Foundation of China (Grant No. 32341008), Shanghai Shuguang Scholars Project, Shanghai Excellent Academic Leader Project, Shanghai Science and Technology Innovation Action Plan-Key Specialization in Computational Biology and Fundamental Research Funds for the Central Universities, and Shanghai Municipal Science and Technology Major Project (Grant No. 2021SHZDZX0100).

## Author Contributions Statement

Qi Liu, Yicheng Gao and Zhiting Wei designed the framework of this work. Yicheng Gao, Zhiting Wei, Kejing Dong, Jingya Yang and Guohui Chuai performed the analyses. Yicheng Gao, Zhiting Wei and Qi Liu wrote the manuscript with the help of other authors. All authors read and approved the final manuscript.

## Competing Interests Statement

The authors declare that they have no competing interests.

**Extended Data Fig. 2:**
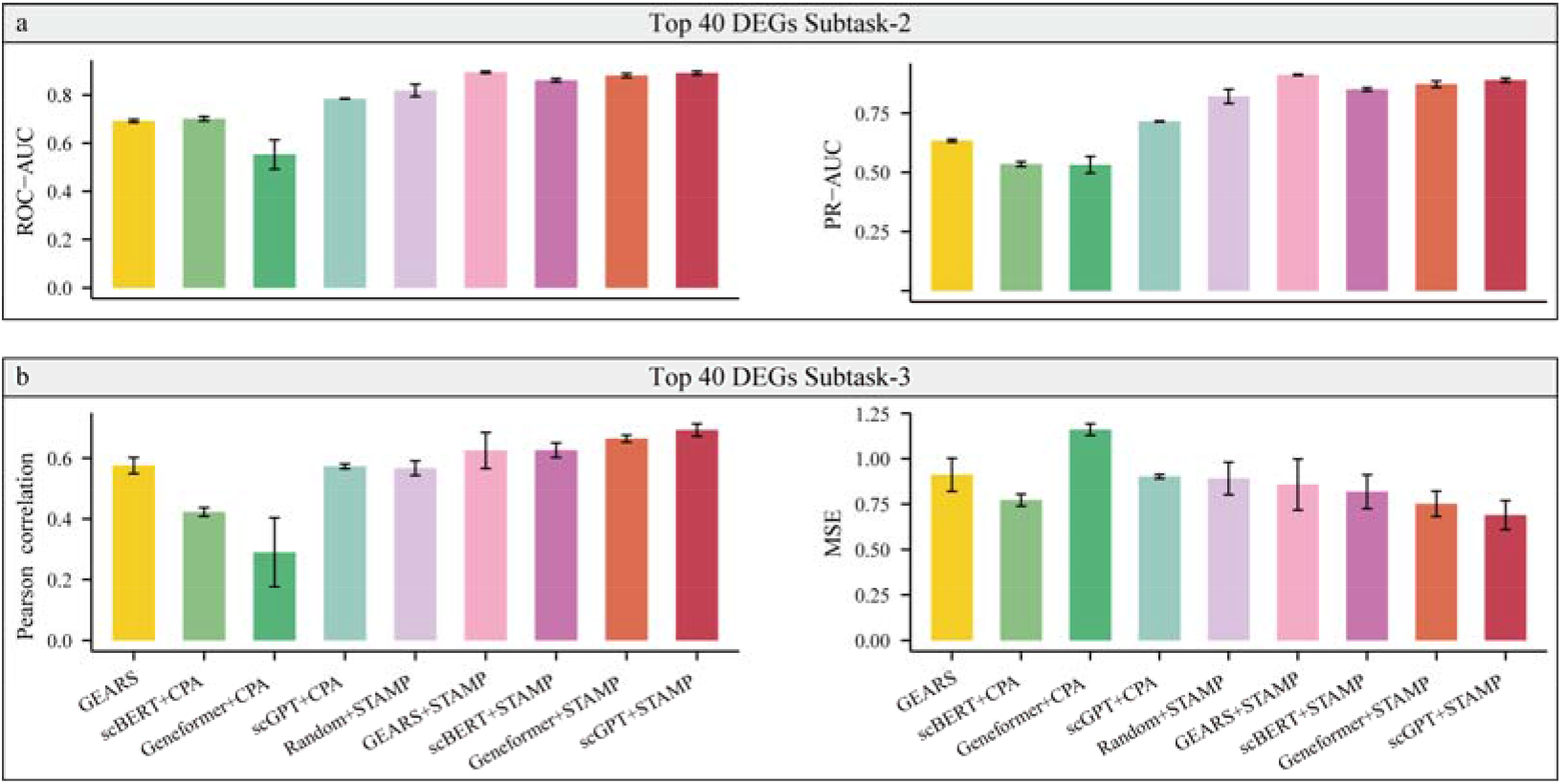
Challenge 1. Single genetic perturbation outcomes predicted for the RPE1_essential dataset (considering the top 40 DEGs). a. The ROC-AUC and PR-AUC values produced by different models in subtask-2. b. The Pearson correlations and MSEs of different models in subtask-3. The data are presented as their mean values and the error bars show the 95% CIs of the mean estimates.

**Extended Data Fig. 4:**
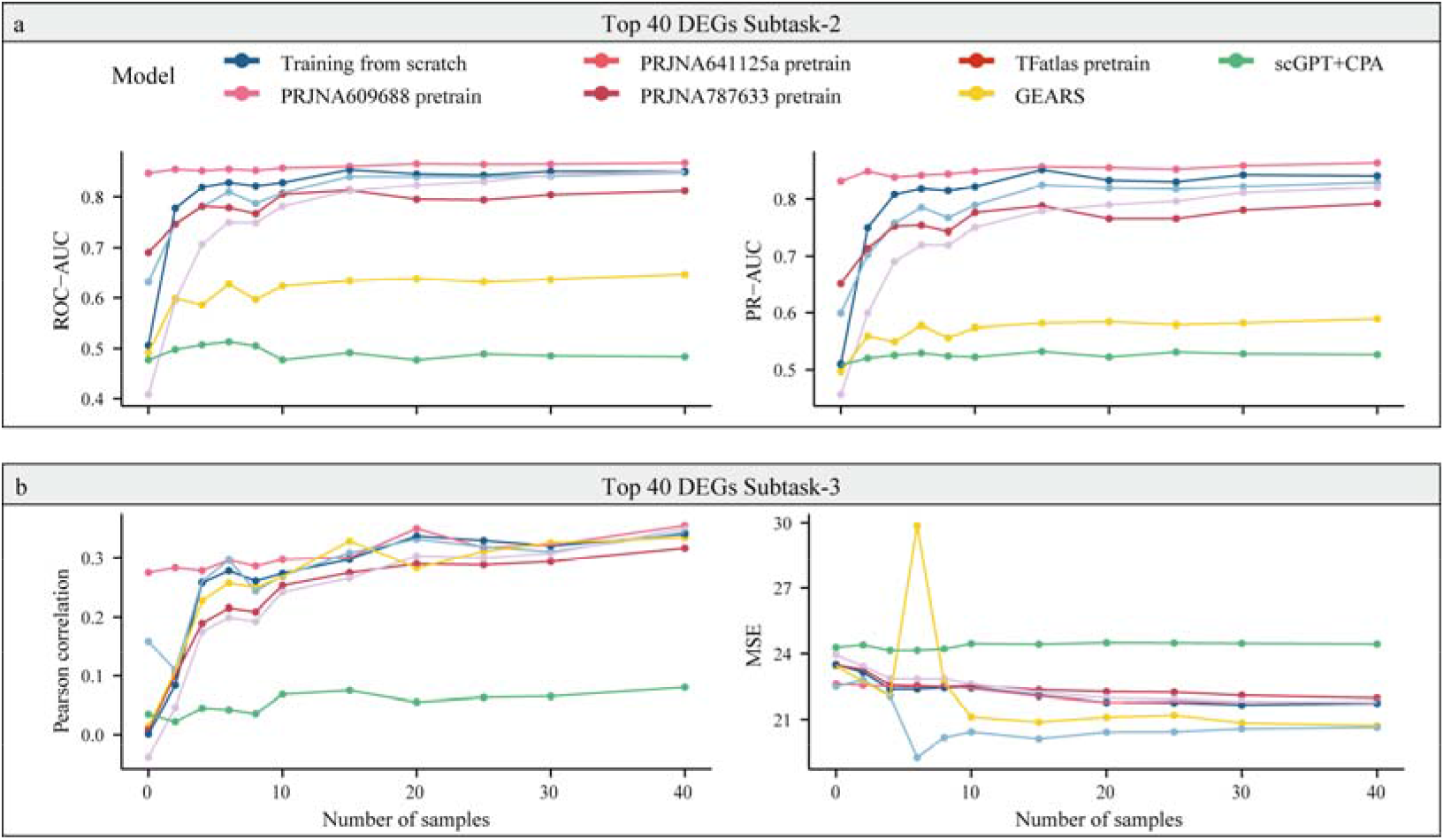
Challenge 3. Genetic perturbation outcomes predicted across cell lines in the PRJNA551220 dataset (considering the top 40 DEGs). a. The ROC-AUC and PR-AUC values produced by different models on pretrained cell line datasets in subtask-2. b. The Pearson correlations and MSEs yielded by different models on pretrained cell line datasets in subtask-3. Each model was fine-tuned with various numbers of samples from the target datasets (0, 2, 4, 6, 8, 10, 15, 20, 25, 30, and 40 samples). The plot shows the average performance as a function of the number of samples acquired from the target cell line dataset.

## References

1. Tang, F. et al. mRNA-Seq whole-transcriptome analysis of a single cell. Nature methods 6, 377–382 (2009).

2. Bock, C. et al. High-content CRISPR screening. Nature Reviews Methods Primers 2, 8 (2022).

3. Dixit, A. et al. Perturb-Seq: dissecting molecular circuits with scalable single-cell RNA profiling of pooled genetic screens. cell 167, 1853–1866. e1817 (2016).

4. Cheng, J. et al. Massively Parallel CRISPR-Based Genetic Perturbation Screening at Single-Cell Resolution. Advanced Science 10, 2204484 (2023).

5. Norman, T.M. et al. Exploring genetic interaction manifolds constructed from rich single-cell phenotypes. Science 365, 786–793 (2019).

6. Kamimoto, K. et al. Dissecting cell identity via network inference and in silico gene perturbation. Nature 614, 742–751 (2023).

7. Qiu, X. et al. Mapping transcriptomic vector fields of single cells. Cell 185, 690–711. e645 (2022).

8. Adamson, B. et al. A multiplexed single-cell CRISPR screening platform enables systematic dissection of the unfolded protein response. Cell 167, 1867–1882. e1821 (2016).

9. Jaitin, D.A. et al. Dissecting immune circuits by linking CRISPR-pooled screens with single-cell RNA-seq. Cell 167, 1883–1896. e1815 (2016).

10. Roohani, Y., Huang, K. & Leskovec, J. Predicting transcriptional outcomes of novel multigene perturbations with gears. Nature Biotechnology, 1–9 (2023).

11. Lotfollahi, M. et al. Predicting cellular responses to complex perturbations in high-throughput screens. Molecular Systems Biology, e11517 (2023).

12. Ji, Y., Lotfollahi, M., Wolf, F.A. & Theis, F.J. Machine learning for perturbational single-cell omics. Cell Systems 12, 522–537 (2021).

13. Aibar, S. et al. SCENIC: single-cell regulatory network inference and clustering. Nature methods 14, 1083–1086 (2017).

14. Hetzel, L., Boehm, S., Kilbertus, N., Günnemann, S. & Theis, F. Predicting cellular responses to novel drug perturbations at a single-cell resolution. Advances in Neural Information Processing Systems 35, 26711–26722 (2022).

15. Inecik, K., Uhlmann, A., Lotfollahi, M. & Theis, F. Multicpa: Multimodal compositional perturbation autoencoder. bioRxiv, 2022.2007. 2008.499049 (2022).

16. Yang, F. et al. scBERT as a large-scale pretrained deep language model for cell type annotation of single-cell RNA-seq data. Nature Machine Intelligence 4, 852–866 (2022).

17. Theodoris, C.V. et al. Transfer learning enables predictions in network biology. Nature, 1–9 (2023).

18. Cui, H. et al. scGPT: Towards building a foundation model for Single-Cell multi-omics using generative AI. bioRxiv, 2023.2004. 2030.538439 (2023).

19. Ma, J. et al. Few-shot learning creates predictive models of drug response that translate from high-throughput screens to individual patients. Nature Cancer 2, 233–244 (2021).

20. Chang, O., Flokas, L., Lipson, H. & Spranger, M. Assessing SATNet ‘s ability to solve the symbol grounding problem. Advances in Neural Information Processing Systems 33, 1428–1439 (2020).

21. Wies, N., Levine, Y. & Shashua, A. in The Eleventh International Conference on Learning Representations (2022).

22. Gülçehre, Ç. & Bengio, Y. Knowledge matters: Importance of prior information for optimization. The Journal of Machine Learning Research 17, 226–257 (2016).

23. Glasmachers, T. in Asian conference on machine learning 17–32 (PMLR, 2017).

24. Wang, P.-W., Donti, P., Wilder, B. & Kolter, Z. in International Conference on Machine Learning 6545-6554 (PMLR, 2019).

25. Zhang, C., Gao, F., Jia, B., Zhu, Y. & Zhu, S.-C. in Proceedings of the IEEE/CVF conference on computer vision and pattern recognition 5317–5327 (2019).

26. Hu, S., Ma, Y., Liu, X., Wei, Y. & Bai, S. in Proceedings of the AAAI Conference on Artificial Intelligence, Vol. 35 1567–1574 (2021).

27. Chollet, F. On the measure of intelligence. arXiv preprint arXiv:1911.01547 (2019).

28. Piękos, P., Malinowski, M. & Michalewski, H. in Proceedings of the 59th Annual Meeting of the Association for Computational Linguistics and the 11th International Joint Conference on Natural Language Processing (Volume 2: Short Papers) 383–394 (2021).

29. Wei, J. et al. Chain-of-thought prompting elicits reasoning in large language models. Advances in Neural Information Processing Systems 35, 24824–24837 (2022).

30. Zhang, Y. & Yang, Q. A survey on multi-task learning. IEEE Transactions on Knowledge and Data Engineering 34, 5586–5609 (2021).

31. Replogle, J.M. et al. Mapping information-rich genotype-phenotype landscapes with genome-scale Perturb-seq. Cell 185, 2559–2575. e2528 (2022).

32. Joung, J. et al. A transcription factor atlas of directed differentiation. Cell 186, 209–229. e226 (2023).

33. Wolf, F.A., Angerer, P. & Theis, F.J. SCANPY: large-scale single-cell gene expression data analysis. Genome biology 19, 1–5 (2018).

34. Squair, J.W. et al. Confronting false discoveries in single-cell differential expression. Nature communications 12, 5692 (2021).

35. Cohen, I. et al. Pearson correlation coefficient. Noise reduction in speech processing, 1–4 (2009).

36. Prasad, N.N. & Rao, J.N. The estimation of the mean squared error of small-area estimators. Journal of the American statistical association, 163–171 (1990).

37. Frangieh, C.J. et al. Multimodal pooled Perturb-CITE-seq screens in patient models define mechanisms of cancer immune evasion. Nature genetics 53, 332–341 (2021).

38. Schmidt, R. et al. CRISPR activation and interference screens decode stimulation responses in primary human T cells. Science 375, eabj4008 (2022).

39. Wang, W., Zheng, V.W., Yu, H. & Miao, C. A survey of zero-shot learning: Settings, methods, and applications. ACM Transactions on Intelligent Systems and Technology (TIST) 10, 1–37 (2019).

40. Gao, Y. et al. Pan-Peptide Meta Learning for T-cell receptor–antigen binding recognition. Nature Machine Intelligence, 1–14 (2023).

41. Hospedales, T., Antoniou, A., Micaelli, P. & Storkey, A. Meta-learning in neural networks: A survey. IEEE transactions on pattern analysis and machine intelligence 44, 5149–5169 (2021).

42. Tian, R. et al. Genome-wide CRISPRi/a screens in human neurons link lysosomal failure to ferroptosis. Nature neuroscience 24, 1020–1034 (2021).

43. Replogle, J.M. et al. Combinatorial single-cell CRISPR screens by direct guide RNA capture and targeted sequencing. Nature biotechnology 38, 954–961 (2020).

44. Wang, Y., Yao, Q., Kwok, J.T. & Ni, L.M. Generalizing from a few examples: A survey on few-shot learning. ACM computing surveys (csur) 53, 1–34 (2020).

45. Tran, H.T.N. et al. A benchmark of batch-effect correction methods for single-cell RNA sequencing data. Genome biology 21, 1–32 (2020).

46. Papalexi, E. et al. Characterizing the molecular regulation of inhibitory immune checkpoints with multimodal single-cell screens. Nature genetics 53, 322–331 (2021).

47. Mimitou, E.P. et al. Multiplexed detection of proteins, transcriptomes, clonotypes and CRISPR perturbations in single cells. Nature methods 16, 409–412 (2019).

48. Young, M.D., Wakefield, M.J., Smyth, G.K. & Oshlack, A. Gene ontology analysis for RNA-seq: accounting for selection bias. Genome biology 11, 1–12 (2010).

49. Moon, J.W. et al. IFNγ induces PD-L1 overexpression by JAK2/STAT1/IRF-1 signaling in EBV-positive gastric carcinoma. Scientific reports 7, 17810 (2017).

50. Garcia-Diaz, A. et al. Interferon receptor signaling pathways regulating PD-L1 and PD-L2 expression. Cell reports 19, 1189–1201 (2017).

51. De Ville, B. Decision trees. Wiley Interdisciplinary Reviews: Computational Statistics 5, 448–455 (2013).

52. Song, B. et al. Decoding Heterogenous Single-cell Perturbation Responses. bioRxiv, 2023.2010. 2030.564796 (2023).

